# Acclimation of phenology relieves leaf longevity constraints in deciduous forests

**DOI:** 10.1101/2022.01.17.476561

**Authors:** Laura Marqués, Koen Hufkens, Christof Bigler, Thomas W. Crowther, Constantin M. Zohner, Benjamin D. Stocker

## Abstract

Leaf phenology is key for regulating total growing season mass and energy fluxes. Long-term temporal trends towards earlier leaf unfolding are observed across Northern Hemisphere forests. Phenological dates also vary between years, whereby end-of-season (EOS) dates correlate positively with start-of-season (SOS) dates and negatively with growing season total net CO_2_ assimilation (*A*_net_). These associations have been interpreted as the effect of a constrained leaf longevity or of premature carbon (C) sink saturation - with far-reaching consequences for long-term phenology projections under climate change and rising CO_2_. Here, we use multi-decadal ground and remote-sensing observations to show that the relationships between *A*_net_ and EOS are opposite at the interannual and the decadal time scales. A decadal trend towards later EOS persists in parallel with a trend towards increasing *A*_net_ - in spite of the negative *A*_net_-EOS relationship at the interannual scale. This indicates that acclimation of phenology has enabled plants to transcend a constrained leaf longevity or premature C sink saturation over the course of several decades, leading to a more effective use of available light and a sustained extension of the vegetation CO_2_ uptake season over time.

## Main Text

For deciduous tree species in temperate and boreal forests, the timing of leaf unfolding in spring and leaf senescence in autumn determines the length of the season during which sunlight is intercepted by leaves, CO_2_ is taken up, and water is transpired. SOS and EOS dates fluctuate at multiple scales, driven by numerous interacting mechanisms that will collectively determine the long-term response to climate change. Dates of leaf phenology vary across climatic^1^ and elevational gradients^2^. Long-term temporal trends towards earlier leaf unfolding have been observed across the Northern Hemisphere in remote sensing data and documented in long-term tree-level observations^3–5^. Such phenological shifts in response to global climate change are altering carbon, water, and nutrient cycling and induce feedbacks within the Earth system^6, 7^.

Relatively reliable models exist to predict SOS based on accumulated temperature and photoperiod^8–11^. In contrast, long-term trends in autumn senescence are less clear^3, 12–14^, depend on EOS definitions based on senescence start, leaf discolouration stages, or dormancy^15, 16^, and drivers are not well understood^6^. Although experimental evidence exists demonstrating that warm autumn temperatures delay leaf senescence^17^, long-term observations often do not show corresponding phenology trends in spite of persistent autumn warming^18, 19^, but see^20^. This has compromised the development of accurate predictive models and undermines phenology projections under future climate conditions^21–23^. However, a positive correlation between annual SOS and EOS dates has been found in observational^24, 25^ and experimental studies^17, 26^, potentially providing useful information for improving EOS predictions. A recent study^27^ found an even stronger relationship between observed EOS and simulated *A*_net_, such that greater productivity was associated with earlier leaf senescence. This negative relationship between *A*_net_ and EOS was interpreted as an expression of plant C sink saturation^26, 28, 29^, whereby an early replenishment of non-structural carbon reserves induces an early cessation of the photosynthetically active season. An EOS advancement over the second half of the 21^st^ century was thus predicted as a consequence of accelerated C sink saturation due to continued SOS advances and enhanced photosynthesis under rising CO_2_ levels^27^. However, in the past, a sustained SOS advance^3^ and a widely observed CO_2_-driven increase in photosynthesis^30–33^ did not lead to a corresponding EOS advance^3^. With these interacting mechanisms operating over different spatio-temporal scales^34^, and the influence of their associated environmental controls, it has been challenging to identify general trends in the changes in autumn leaf senescence.

Here, we investigated this apparent conflict by decomposing long-term trends, interannual, and spatial variations using linear mixed-effects models (LMMs). We hypothesized that the relationships between *A*_net_ and EOS are driven by multiple processes and are non-stationary over decadal time scales. We complemented the analysis of multi-year ground observations (1948-2015) from the PEP725 dataset^5^ of 434,226 European tree-level phenology observations with an analysis of remotely sensed phenological dates to expand the extent of data coverage in spatial and climatic space. Remotely sensed estimates of phenology (2001-2018) were obtained from MODIS MCD12Q2 Collection 6^35, 36^ for 4,879 randomly sampled points of deciduous tree species in temperate and boreal forests in the northern hemisphere. We explored the robustness of *A*_net_ estimates and respective statistical models by using *A*_net_ estimates as previously used by *Zani et al*.^27^ and performed all analyses also with estimates generated here using an alternative, comprehensively evaluated photosynthesis model^37^. See Methods for a detailed account of the analysis, data, and modelling.

We found opposing *A*_net_-EOS relationships at different temporal scales. When controlling for the effect of *A*_net_, we found a clear decadal-scale trend component towards later EOS (0.253 ± 0.001 d yr^-1^, Fig. 1A). After separating the long-term trend, the remaining *A*_net_-EOS relationship reflects interannual variations (Fig. 1B). At this scale, *A*_net_ is negatively correlated with EOS, as reported by *Zani et al*.^27^ based on a univariate model (Fig. 1C). The net effect of these opposing relationships is a relatively small delay in EOS over time (0.046 ± 0.001 d yr^-1^, fig. S1A) - in spite of the steadily increasing (simulated) *A*_net_ since the mid-20^th^ century (fig. S1B). These results are robust against the use of alternative *A*_net_ estimates^37, 38^ in LMMs (figs. S1C, S2A and S2B). The parallel gradual *A*_net_ increase and long-term trend towards delayed autumn senescence indicate a positive *A*_net_-EOS relationship - opposite to the negative relationship at the interannual time scale.

**Fig. 1.**
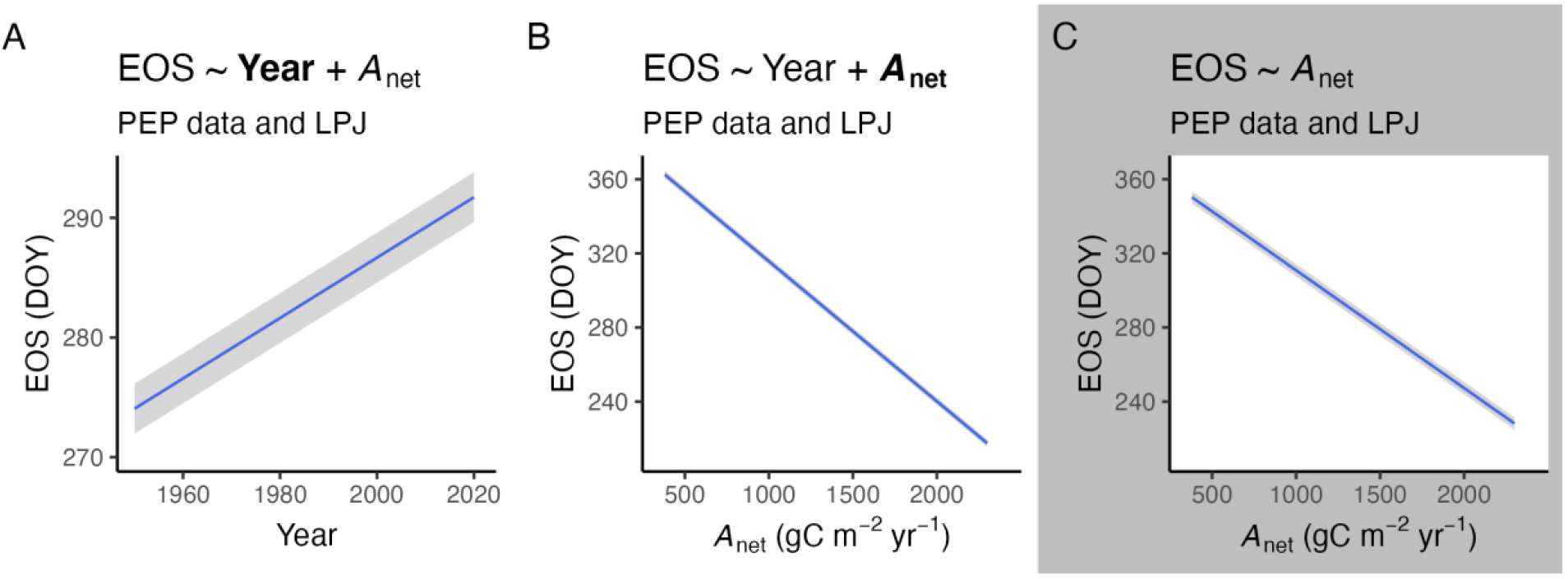
Relationship of CO_2_ assimilation and autumn phenology from local observations (PEP725 data). (**A, B**) Partial relationships of a multiple LMM, where end-of-season (EOS, expressed as day-of-year, DOY) is the response variable and (**A**) the long-term trend (year) and (**B**) *A*_net_ (simulated using the LPJ model) are treated as fixed effects. (**C**) EOS versus *A*_net_ based on an LMM with *A*_net_ as a single fixed effect. Blue lines represent the expected values from LMMs and grey ranges their 95% confidence intervals. In both bivariate and univariate models, site and species are treated as grouping variables of random intercepts.

Scale-dependent relationship reversals were found also when decomposing interannual from spatial variations, i.e. when separating annual anomalies from multi-year means by site in LMMs. Due to relatively limited temporal coverage of the remote sensing data (2001-2018), we did not separate a long-term trend. Across space, higher mean *A*_net_ is associated with later mean EOS (Fig. 2A), while the opposite relationship prevails when considering interannual variations at a given location (Fig. 2B). The positive mean EOS-mean *A*_net_ relationship is also evident when considering the spatial distribution of observations across the Northern Hemisphere (Figs. 2C and 2D).

**Fig. 2.**
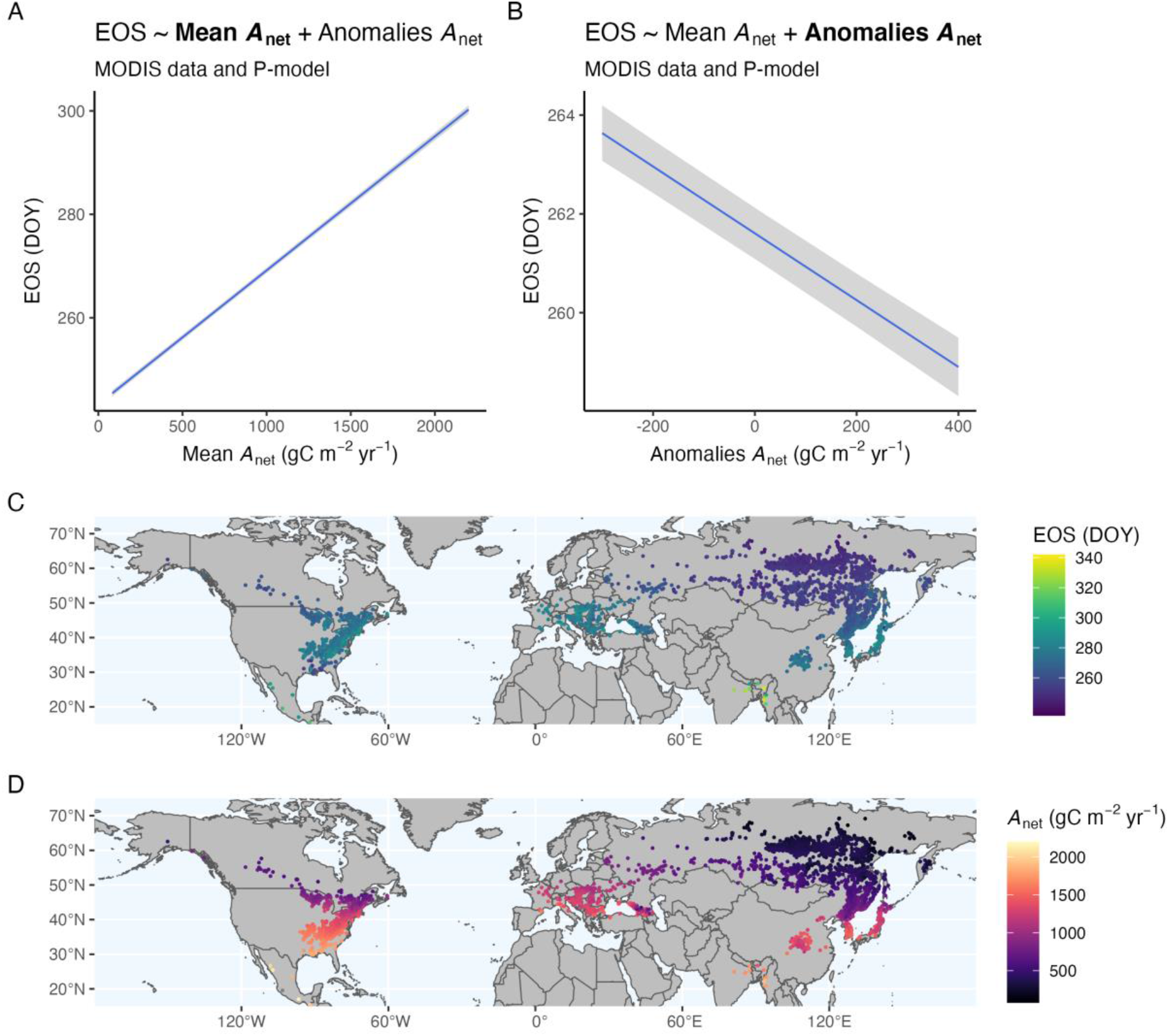
Relationships of CO_2_ assimilation and autumn phenology from remote-sensing observations (MODIS C6 MCD12Q2 data). (**A, B**) Partial relationships of a multiple LMM, with *A*_net_ simulated using the P-model, and where both (**A**) *A*_net_ mean and (**B**) anomalies relative to the mean value from 2001 to 2018 are treated as fixed effects, and site and year are treated as grouping variables of random intercepts. Blue lines represent the expected values from LMMs and grey ranges their 95% confidence intervals. (**C, D**) Mean values of (**C**) EOS and (**D**) *A*_net_ simulated by the P-model for grid cells distributed along the Northern Hemisphere.

These relationships yield several insights into potential processes underlying phenology shifts under global environmental change. A link between interannual variations of *A*_net_ and EOS emerges from both the ground-based and remote sensing-based analyses. Since *A*_net_ represents the cumulative net CO_2_ assimilation since SOS, the *A*_net_-EOS relationship is closely related to the previously reported relationship between SOS and EOS^17, 24–26^. Early leaf unfolding leading to early senescence has been hypothesized to be the result of a relatively constant length of leaf phenological stages^39^, or of a leaf aging effect^40–42^, whereby a tightly constrained leaf longevity implies direct control of SOS on EOS^24^. Given the well-documented gradual SOS advancement^43–47^ (fig. S1D), this process should induce an advancement also of EOS. Our analysis reveals that this has not been the case (figs. S3 and S4). Similarly, also the strong relationship between *A*_net_-EOS, apparent at the interannual scale, has not been stationary over several decades. This indicates that the interplay between multiple drivers and processes has resulted in a gradual relief of tight constraints relevant at the interannual time scale, potentially arising from premature leaf aging or C sink saturation. We interpret this non-stationarity of the *A*_net_-EOS and the SOS-EOS relationships as being reflective of acclimation.

What are the drivers of observed phenological relationships and their acclimation? If the negative interannual *A*_net_-EOS relationship was due to C sink saturation, it should prevail also in the long-term, as photosynthetic CO_2_ assimilation is enhanced under rising atmospheric CO_2_^30–33^. However, it appears that the tight constraints, apparent at the interannual scale, are relieved over the course of several decades. The opposing relationships at different scales found here question the prediction that gradual increases in photosynthesis cause a progressive advancement of EOS by earlier C sink saturation. Previous studies have generally demonstrated a thermal control delaying EOS across spatial gradients and years^48^. Rising autumn temperatures, causing a slow-down of chlorophyll degradation and leaf discoloration^15^, have been suggested to underlie trends and are considered in autumn senescence models^49^. Most warming experiments have also shown later EOS for various deciduous tree species^50^. Further insights into the importance of a potential C sink saturation mechanism causing a reversal of effects by warming autumn temperatures will be gained by linking observations of non-structural C dynamics with autumn phenology. Our results indicate that different mechanisms and environmental controls are at play at different temporal scales, potentially undermining long-term projections, informed by short-term observations of autumn phenology.

Why does a negative relationship between *A*_net_ and EOS emerge when the long-term trend and the spatial variation are not separated in LMMs? A possible explanation is that relatively large interannual phenology variations dominate over the smaller long-term temporal pattern in the data we analysed, and mask their effect in univariate models. Indeed, separating the opposing long-term and spatial trends from the remaining component of interannual variations improves model performance significantly (ANOVA p < 0.001 and lower AIC) and increases the strength of their interannual links in all models (see estimates for bivariate and univariate models in table S1). This provides further support for an important mechanism underlying the apparent link at the interannual scale. However, it also indicates that this does not preclude the existence of other mechanisms, enabling an acclimating response, and leading to a relief of phenology relationships apparent at the interannual scale and to a sustained delay of EOS. The net effect of opposing mechanisms, apparent in the univariate models, is subject to the data and their relative magnitudes of variations across multiple scales.

The long-term and spatial relationships between *A*_net_ and EOS (and between SOS and EOS) are qualitatively consistent. This indicates that the phenology of individual trees has acclimated over the course of decades in the same direction as evident from the spatial analysis (high *A*_net_ occurs in places with late EOS, Figs. 2C and D). Note that the long-term temporal trends derived from tree-level observations in the PEP data emerge within individual species observed at different locations, while spatial variations may arise also as a result of varying species composition across space and of adaptation within populations of a given species. Our results suggest a clear plasticity of autumn phenology over time, mirroring effects by species distribution and long-term adaptation of individuals and plant communities growing along a large climatic gradient. This indicates a trend towards optimal, climate-adapted functioning.

Opposing relationships at different time scales and across the Northern Hemisphere reconcile apparent conflicts by the reported negative *A*_net_-EOS relationship but absent shifts towards earlier EOS as *A*_net_ has increased over past decades. We conclude that a gradual acclimation of plant physiology and adaptation of phenology has enabled plants to transcend a constrained leaf longevity or a stationary C sink saturation effect, evident from the clear short-term SOS-EOS and *A*_net_-EOS relationships. Thus, in the long run, plants may assimilate more CO_2_ without a direct and inescapable penalty by earlier leaf senescence and without thus foregoing late-season carbon assimilation. This apparent plasticity in phenology appears to have driven plants towards optimal functioning in a changing climate.

## Methods

### Pan European observational data

Spring and autumn phenology dates were collected from the Pan European Phenology Project^5^, which provides in-situ observations for Europe. Phenology dates were defined following the BBCH (Biologische Bundesanstalt, Bundessortenamt und Chemische Industrie) codes and the data selection made by *Zani et al*.^27^. Leaf-out was defined as the date when the first (BBCH11) or 50% of leaf stalks are visible (BBCH13) for the deciduous angiosperms, and as the date when the first leaves separated (BBCH10) for the deciduous conifers. Leaf senescence was defined as the date when 50% of leaves had lost their green color (BBCH94) or had fallen (BCCH95). Following *Zani et al*.^27^ data cleaning, the dataset resulted in 3,855 sites across Central Europe with 14,626 individual time series and 434,226 phenological observations between 1948 and 2015.

### MODIS phenology data

We used the MODIS C6 MCD12Q2 Land Surface Dynamics Product^35^ which provides land surface phenological data at 500-meter spatial resolution from 2001 to 2018, derived from time series of the 2-band Enhanced Vegetation Index (EVI2) calculated from MODIS Nadir Bi-directional Reflectance Distribution Function (BRDF) adjusted surface reflectance (NBAR-EVI2)^36, 51^. From this product, leaf-out (start-of-season, SOS) was taken as the MidGreenup point, i.e., the date when EVI2 first crossed 50% of the segment EVI2 amplitude. Leaf senescence (end-of-season, EOS) was taken as the MidGreendown point, i.e., the date when EVI2 last crossed 50% of the segment EVI2 amplitude. We selected the MidGreenup and MidGreendown to define SOS and EOS instead of the Greenup and Dormancy following the advice from the MCD12Q2 Product user guide to capture the season start and end in high-latitude regions. Data were downloaded using the *MODISTools* R package^52^. We randomly sampled 5,000 pixels spread evenly between temperate deciduous needle and broadleaf IGBP classes and selected the points corresponding to the Northern Hemisphere (4,879).

### Photosynthesis estimation

For locations where tree-level phenology observations were available from the PEP data, we used two alternative estimates of *A*_net_. Results shown in Fig. 1 are based on *A*_net_ as estimated by *Zani et al*.^27^ using their implementation of the LPJ model^38^ and represents gross assimilation minus daytime dark respiration. The cumulative growing season net photosynthesis (*A*_net_) was then obtained by summing the daily *A*_net_ for all days of the growing season, starting at the date of observed SOS as given by the PEP data and ending on the date when daylength falls below 11.2 hours. A detailed explanation of the seasonal photosynthesis estimation is provided in *Zani et al*.^27^. Results shown in figs. S1C and S2 are based on estimates of *A*_net_ using the P-model^37^ as implemented by the *rsofun* R package^53^, and represent gross assimilation minus dark respiration. Cumulative *A*_net_ was calculated from days starting at the observed SOS and ending at the summer solstice (21^st^ of June in the Northern Hemisphere). The same P-model based approach was applied for estimating cumulative *A*_net_ at locations where phenology data was extracted from the MODIS remote sensing product. The P-model predicts leaf-level acclimation of photosynthesis to its environment and simulates CO_2_ assimilation as a linear function of absorbed photosynthetically active radiation (APAR). Here, APAR is based on shortwave radiation from WATCH-WFDEI^54^ and downscaled using WorldClim2^55^, assuming a fraction of APAR of 1.0 for all sites and dates between their respectively observed SOS and EOS dates. Hence, *A*_net_ represents a leaf-level quantity, representative for conditions in full light. Also other meteorological forcing data for P-model simulations were taken from WATCH-WFDEI^54^, downscaled using WorldClim2^55^ as implemented by the *ingestr* R package^56^. Details of the theory underlying the P-model are described in *Stocker et al*.^37^) and *Wang et al*.^57^.

### Data analysis

We fitted linear mixed-effects models to investigate the relationships between autumn phenology (EOS), net photosynthesis (*A*_net_) and spring leaf-out (SOS). We were particularly interested in separating the interannual, the long-term, and the spatial components of variation. The general structure of the models can be summarized as:

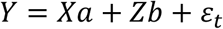

where *Y* represents the dependent variable (i.e., EOS, expressed as day-of-year), *a* is the vector of fixed effects, *b* is the vector of random intercepts, *X* and *Z* are regression matrices of fixed and random effects, respectively, and *ε*_*t*_ is the within-group error vector. For performing the temporal analyses, the predictor variables (*A*_net_, SOS, year) were standardized and site and species were treated as grouping variables of random intercepts. For performing the spatial analysis, we calculated the mean and standard deviation values of the predictors (*A*_net_ and SOS) and evaluated their effects on EOS. Site and year were treated as grouping variables of random intercepts for spatial analyses with MODIS data. Residuals of the models were checked for normality and homoscedasticity. Linear mixed-effects models were fitted using the *lme4* R package^58^. The analyses were performed using the *R* statistical software version 4.0.5^59^.

## Acknowledgments

We thank Trevor Keenan and Yann Vitasse for helpful comments on the manuscript. LM and BDS were funded by the Swiss National Science Foundation grant no. PCEFP2_181115. KH was supported by the generosity of Eric and Wendy Schmidt by recommendation of the Schmidt Futures program. CMZ was funded by the Ambizione grant PZ00P3_193646. Data were provided by the members of the PEP725 project.

## Author contributions

BDS and LM conceived and developed the study; CMZ gathered the PEP data and ran the LPJ simulations; BDS, KH and LM gathered the MODIS data, ran the P-model simulations, and conducted the statistical analyses; LM and BDS led the writing of the manuscript; CB contributed critically to the analyses and the writing; CMZ and TC gave substantial inputs to the manuscript. All authors gave final approval for publication.

## Competing interests

The authors declare no competing interests.

## Data and materials availability

Code for the data analysis of this study is available at the Github repository DOI:10.5281/zenodo.5799643. Ground phenology data provided by the members of the PEP725 project is freely available at http://www.pep725.eu. Remote-sensing phenology data from the MODIS C6 MCD12Q2 Land Surface Dynamics Product is freely accessible at https://lpdaac.usgs.gov/products/mcd12q2v006/.

## Notes

### Competing Interest Statement

The authors have declared no competing interest.

