## Supplementary material for "Acclimation of phenology relieves leaf longevity constraints in deciduous forests"

**Content**

Supplementary Figs. S1 to S4
Supplementary Table S1

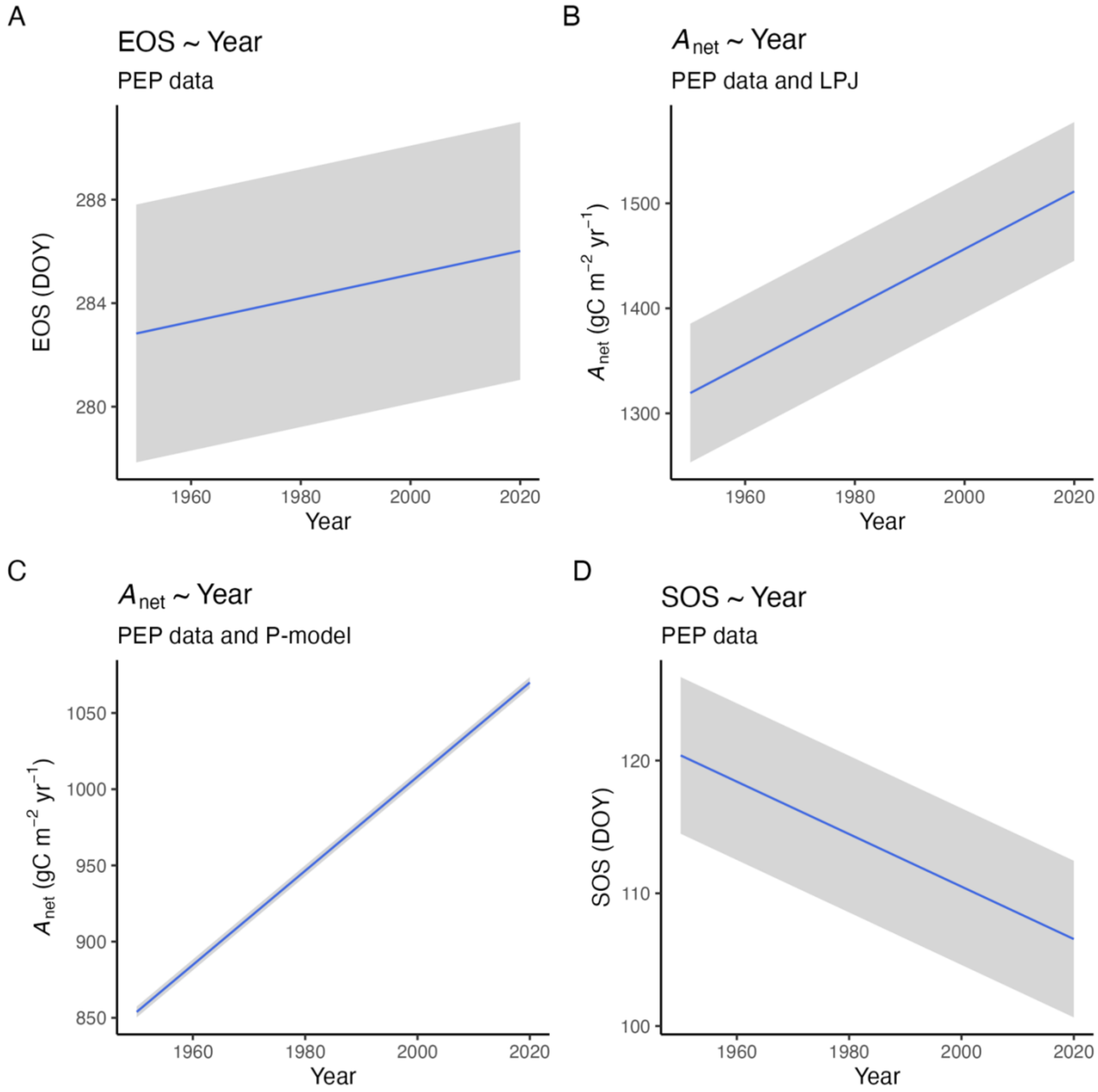


**Fig. S1. Temporal trends of $CO_2$ assimilation and phenological dates from local observations (PEP725 data).** (**A**) Trends towards later EOS, (**B**) higher $A_{net}$ from LPJ simulations and (**C**) from P-model simulations, and (**D**) earlier SOS based on an LMM with year as a single fixed effect and site and species as grouping variables of the random intercepts. Blue lines represent the expected values from LMMs and grey ranges their 95% confidence intervals.

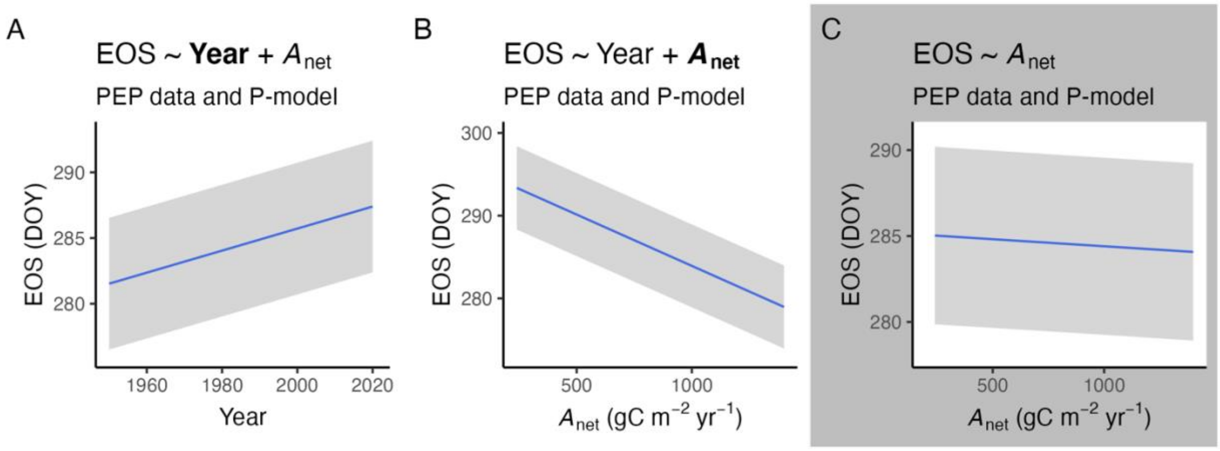


**Fig. S2. Relationship of $CO_2$ assimilation and autumn phenology from local observations (PEP725 data).** (**A, B**) Partial relationships of a multiple LMM, where end-of-season (EOS, expressed as day-of-year, DOY) is the response variable and (**A**) the long-term trend (year) and (**B**) $A_{net}$ (simulated using the P-model) are treated as fixed effects. (**C**) EOS versus $A_{net}$ based on an LMM with $A_{net}$ as a single fixed effect. Blue lines represent the expected values from LMMs and grey ranges their 95% confidence intervals. In both bivariate and univariate models, site and species are treated as grouping variables of random intercepts.

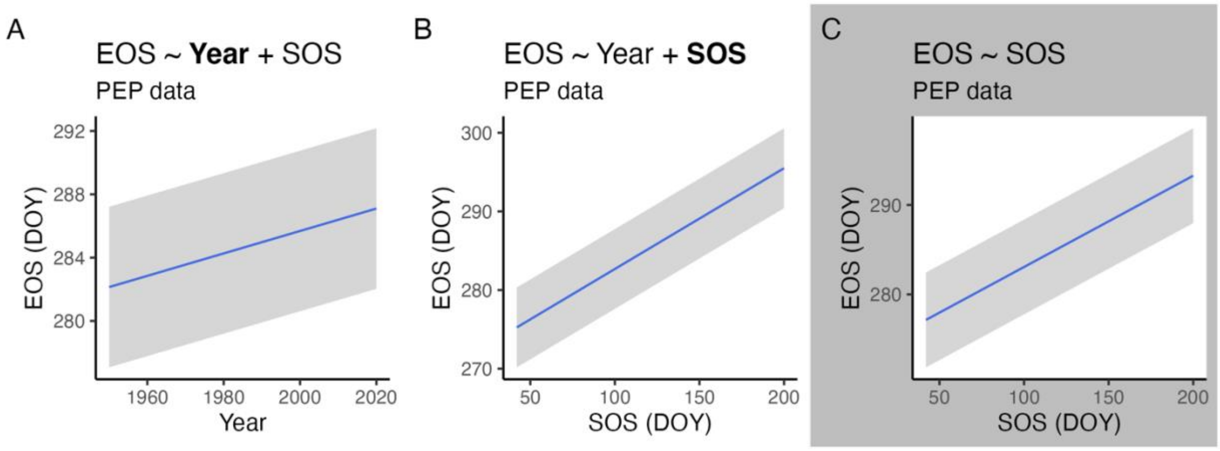


**Fig. S3. Relationship of spring and autumn phenological dates from local observations (PEP725 data).** (**A, B**) Partial relationships of a multiple LMM, where end-of-season (EOS, expressed as day-of-year, DOY) is the response variable and (**A**) the long-term trend (year) and (**B**) SOS are treated as fixed effects. (**C**) EOS versus SOS based on an LMM with SOS as a single fixed effect. Blue lines represent the expected values from LMMs and grey ranges their 95% confidence intervals. In both bivariate and univariate models, site and species are treated as grouping variables of random intercepts.

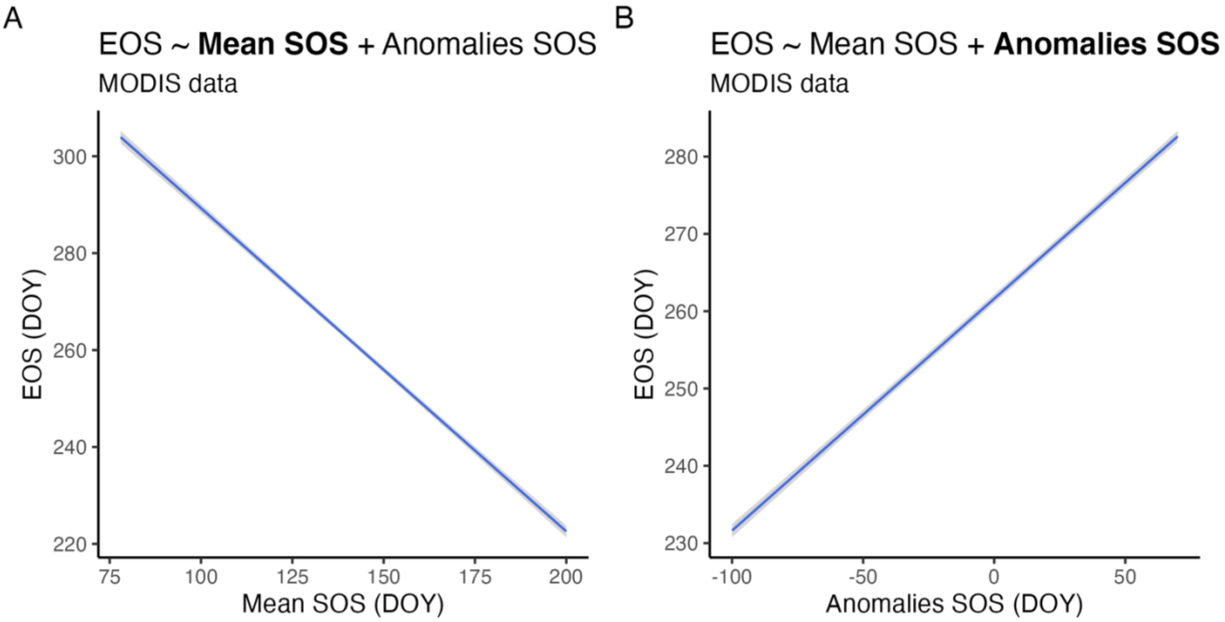


**Fig. S4. Relationships of phenological dates from remote-sensing observations (MODIS C6 MCD12Q2 data).** (**A, B**) Partial relationships of a multiple LMM where both (**A**) SOS mean and (**B**) anomalies relative to the mean value from 2001 to 2018 are treated as fixed effects, and site and year are treated as grouping variables of random intercepts. Blue lines represent the expected values from LMMs and grey ranges their 95% confidence intervals.

**Table S1. Main statistics of the bivariate and univariate LMMs.** Models were fitted to describe the relationships of $CO_2$ assimilation and phenological dates from local observations and remote-sensing estimates. Variables were standardized so effect sizes are directly comparable. $\sigma_u$ denotes the standard deviation of the random intercepts.

| **Model / Figure** | **Response variable** | **Predictor** | **Estimate** | **Standard Error** | ***P-value*** | **Random effects ($\sigma_u$)** | **Data** |
|---|---|---|---|---|---|---|---|
| Bivariate 1A - B | EOS | (Intercept) | 282.823 | 1.064 | <0.001 | Site (9.614) | PEP725 |
| | | Year | 4.139 | 0.018 | <0.001 | Species (2.579) | |
| | | $A_{net}$ (LPJ) | -10.451 | 0.020 | <0.001 | | |
| Univariate 1C | EOS | (Intercept) | 283.167 | 1.701 | <0.001 | Site (9.011) | PEP725 |
| | | $A_{net}$ (LPJ) | -8.797 | 0.019 | <0.001 | Species (4.152) | |
| Bivariate 2A - B | EOS | (Intercept) | 261.606 | 0.264 | <0.001 | Site (6.113) | MODIS |
| | | Mean $A_{net}$ (P-model) | 10.823 | 0.090 | <0.001 | Year (1.049) | |
| | | Anomalies $A_{net}$ (P-model) | -0.442 | 0.024 | <0.001 | | |
| Univariate | EOS | (Intercept) | 261.614 | 0.123 | <0.001 | Site (8.498) | MODIS |
| | | $A_{net}$ (P-model) | 4.976 | 0.093 | <0.001 | | |
| Bivariate S2A - B | EOS | (Intercept) | 284.441 | 2.557 | <0.001 | Site (6.605) | PEP725 |
| | | Year | 1.375 | 0.029 | <0.001 | Species (6.257) | |
| | | $A_{net}$ (P-model) | -1.023 | 0.030 | <0.001 | | |
| Univariate S2C | EOS | (Intercept) | 284.438 | 2.630 | <0.001 | Site (6.539) | PEP725 |
| | | $A_{net}$ (P-model) | -0.068 | 0.023 | 0.003 | Species (6.435) | |
| Bivariate S3A - B | EOS | (Intercept) | 284.602 | 2.587 | <0.001 | Site (6.560) | PEP725 |
| | | Year | 1.162 | 0.022 | <0.001 | Species (6.330) | |
| | | SOS | 1.654 | 0.024 | <0.001 | | |
| Univariate S3C | EOS | (Intercept) | 284.598 | 2.702 | <0.001 | Site (6.600) | PEP725 |
| | | SOS | 1.317 | 0.023 | <0.001 | Species (6.613) | |
| Bivariate S4A - B | EOS | (Intercept) | 261.615 | 0.276 | <0.001 | Site (8.690) | MODIS |
| | | Mean SOS | -8.906 | 0.126 | <0.001 | Year (1.044) | |
| | | Anomalies SOS | 1.876 | 0.021 | <0.001 | | |
| Univariate | EOS | (Intercept) | 261.622 | 0.216 | <0.001 | Site (14.993) | MODIS |
| | | SOS | 3.647 | 0.047 | <0.001 | | |
